# Spatial distribution and functional integration of displaced ipRGCs

**DOI:** 10.1101/2023.09.05.556383

**Authors:** Sabrina Duda, Christoph T. Block, Dipti R. Pradhan, Yousef Arzhangnia, Martin Greschner, Christian Puller

**Affiliations:** Visual Neuroscience, Department of Neuroscience, Carl von Ossietzky University, 26111 Oldenburg, Germany

## Abstract

The mammalian retina contains many distinct types of ganglion cells, which form mosaics to evenly tile the retina with cells of each type at each position of the visual field. It is well known that displaced retinal ganglion cells (dRGCs) exist with cell bodies in the inner nuclear layer, along with regularly placed RGCs with cell bodies in the ganglion cell layer. A prominent example of dRGCs are M1-type intrinsically photosensitive ganglion cells (ipRGCs) which exist in various species including humans and non-human primates. Little is known, however, about their spatial relationship with regularly placed ipRGCs.

Here, we identified mouse ipRGC types M1, M2, and M4/sONɑ by immunohistochemistry and light microscopy to anatomically investigate the distribution of displaced and regularly placed cells. Reconstruction of immunolabeled dendritic mosaics from M1 and sONɑ RGCs indicated that dRGCs tiled the retina evenly with their regularly placed RGC partners. Multi-electrode array recordings revealed conventional receptive fields of displaced sONɑ RGCs which fit into the functional mosaic of their regularly placed counterparts. We further analyzed the RGC distributions across complete retinas. The analysis of regularly placed M1 ipRGCs and ɑRGCs revealed distinct density gradients where ∼16% and ∼8% occurred as dRGCs, respectively. The density distributions of dRGCs showed type-specific patterns which followed neither the global density distribution of all ganglion cells nor the local densities of corresponding cell types.

Our study shows that the displacement of ganglion cell bodies into the inner nuclear layer occurs in a type-dependent manner, where dRGCs are positioned to form complete mosaics with their regularly placed RGC partners. Our data suggest that dRGCs and regularly placed RGCs serve the same functional role within their corresponding population of ganglion cells.

**Significance statement:** We applied large-scale anatomical and electrophysiological experiments in mice to show that displaced intrinsically photosensitive retinal ganglion cells (ipRGCs) complete the mosaics of their regularly placed counterparts with their dendritic trees and receptive fields. Therefore, displaced ipRGCs likely serve the same functional role as corresponding regularly placed cells. The density distributions of displaced ipRGCs showed distinct, type-specific patterns. Interestingly, they followed neither the global density distribution of all ganglion cells nor the local densities of corresponding cell types.

## Introduction

Retinal ganglion cells (RGCs) consist of a heterogeneous group of more than 40 different cell types in mice (Baden et al., 2016; Tran et al., 2019). They can be separated based on the large diversity of morphologies, molecular identities, and light response properties (Baden et al., 2016; Doi et al., 1995; Sun et al., 2002; Tran et al., 2019). Cell bodies of individual RGC types are regularly spaced and their dendrites are arranged in a mosaic which tiles the retina without leaving gaps (Wässle et al., 1981; Wässle and Riemann, 1978). Regional specializations of these mosaics exist across the mammalian retina in response to lifestyle and habitat of different species, such as the cat area centralis, the rabbit visual streak, or the human fovea (Curcio and Allen, 1990; Oyster et al., 1981; Stone, 1965). These regions contain increased RGC densities to enhance visual acuity, for instance, in distinct areas of the visual field (Bleckert et al., 2014; Collin, 2008). In mice, RGCs have been thought to have a central to peripheral density gradient (Dräger and Olsen, 1981). However, recent work showed distinct density gradients for certain types, such as the sustained ON alpha (sONɑ) RGC, contrary to the overall density distribution of RGCs (Salinas-Navarro et al., 2009a; Zhang et al., 2012; Bleckert et al., 2014; Rousso et al., 2016; reviewed in Heukamp et al., 2020).

Independent of these specializations, a common feature of all RGC types is their cell body position in the ganglion cell layer (GCL). However, there are poorly understood exceptions to this rule, where cell bodies of displaced RGC (dRGCs) are located in the inner nuclear layer (INL). A well-known example RGC group which frequently exhibits displaced cell bodies are intrinsically photosensitive ganglion cells (ipRGCs) in various mammalian retinas including human (e.g. (Provencio et al., 2000; Berson et al., 2002; Hattar et al., 2002; Berson et al., 2010; Nasir-Ahmad et al., 2019; Haverkamp et al., 2022). ipRGCs express the photopigment melanopsin and mediate distinct functions in image- and non-image forming vision (reviewed in Aranda and Schmidt, 2021). In the mouse retina, six different types of ipRGCs (M1-M6) can be separated based on morphology, functional properties, and axonal projections to distinct brain regions (Berson et al., 2002; Ecker, 2010; Quattrochi et al., 2019; Schmidt et al., 2011, 2014; Stabio et al., 2018; Berry et al., 2023). The M4 ipRGC was shown to correspond to one of the four alpha RGC types of the mouse retina, i.e. the sustained ON alpha (sONɑ) RGC (Estevez et al., 2012; Schmidt et al., 2014).

Despite the knowledge of this heterogeneous group of ipRGCs and the functional roles of corresponding types, little is known about the properties of displaced ipRGCs. In mice, about 2% of all RGCs are located as displaced retinal ganglion cells (dRGCs) in the inner nuclear layer (INL). They occur more frequently in the retinal periphery (Dräger and Olsen, 1981). Individual dRGCs show a large morphological variety (Pang and Wu, 2011) but dRGCs have not been comprehensively examined with regard to cell identity and regularly placed RGCs. Thus, the spatial arrangement of their dendritic processes with respect to regular RGCs in the GCL or their response properties remain unknown. While dRGCs are often ignored when RGC mosaics are investigated in the mammalian retina, it is thought that dRGCs are misplaced due to ontogenetic aberrations rather than representing an independent functional class of RGCs (Buhl and Dann, 1988; Doi et al., 1995). In birds, however, dRGCs mostly project to the accessory optic system and presumably support the optokinetic nystagmus and retinal image stabilization (Cook and Podugolnikova, 2001; Dann and Buhl, 1987; Simpson, 1984).

To elucidate the anatomical and functional arrangement of dRGCs relative to mosaics of regular RGCs in a mammalian retina, we took advantage of available immunomarkers against the group of ipRGCs in mice. We analyzed the spatial distribution and dendritic arrangement of the ipRGC types M1, M2, and sONɑ, in both GCL and INL of the retina. dRGCs completed the dendritic mosaic and tiled the retina evenly with their regularly placed partners to form complete anatomical and functional mosaics. In addition, we found that the distribution of identified dRGCs neither followed the overall density distribution of all RGCs, nor the distribution of the corresponding type in the GCL. Furthermore, the proportion of displaced cells differed between types. Our data suggest that a displacement of cell bodies does not occur randomly but in a type-dependent manner.

## Material and Methods

### Animals and tissue preparation

All experiments were performed in accordance with the institutional guidelines for animal welfare and the laws on animal experimentation issued by the EU and the German government. Wild type C57Bl6/J mice and *thy1-*GFP-O mice (Feng et al., 2000) of either sex were included in this study. They were housed in a 12:12 hour light dark cycle. Fully grown C57Bl6/J animals at an age of 11-12 weeks were used for cell quantifications. The animals were deeply anesthetized by carbon dioxide followed by a decapitation. Eyes were enucleated and fixed in 4% cold paraformaldehyde in 0.01M phosphate buffered saline (PBS), pH 7.4, for 15 minutes at room temperature. After washing in PBS, special care was taken to maintain the orientation of the eyecups, where the choroid fissure was used as a landmark. Four radial relieving cuts were made, the longest one along the nasal fissure, approaching the optic nerve head. Then, the retinas were dissected away from the eye cup in PBS. This was performed in a way that the complete retina, including the most peripheral region (outer marginal zone), was preserved. Retinas were then cryoprotected overnight with 30% sucrose in PBS, and stored at -20°C until use.

*thy1-*GFP-O mice at an age of 5-7 weeks were used for multi-electrode array (MEA) recordings. Animals were dark adapted for 2 hours and killed by cervical dislocation. Dorsal-temporal regions of the retinas were dissected from the eye cup under infrared illumination in Ames’ solution, pH 7.4 (USBiological) bubbled with carbogen (95% O_2_ and 5% CO_2_) at room temperature. MEA recordings were performed at 36.5 °C in the recording chamber.

### Immunohistochemistry

Following dissection, retinas were used as whole mounts. Immunohistochemical labeling was performed by an indirect immunofluorescence method. The tissue was mounted on a black nitrocellulose filter membrane (Millipore) with GCL up and pre-incubated at room temperature for 2-4 hours in an incubation solution containing 5% NDS, 1% BSA, 1% Triton X-100, and 0.02% sodium azide dissolved in PBS. Primary and secondary antibodies were diluted in the same incubation solution. Wholemounts were incubated with primary antibodies (table 1) for 3 days at room temperature. Secondary antibodies (Alexa 488 and Alexa 568, Invitrogen, 1:500; Alexa 647, Invitrogen or Jackson ImmunoResearch, 1:250) were incubated at room temperature for 4 hours. 4′,6-Diamidino-2-phenylindole dihydrochloride (DAPI, Sigma-Aldrich, cat# D9542, final concentration 0.2µg/ml) was added into the secondary antibody solution for 1 hour before rinsing.

**Table 1:**
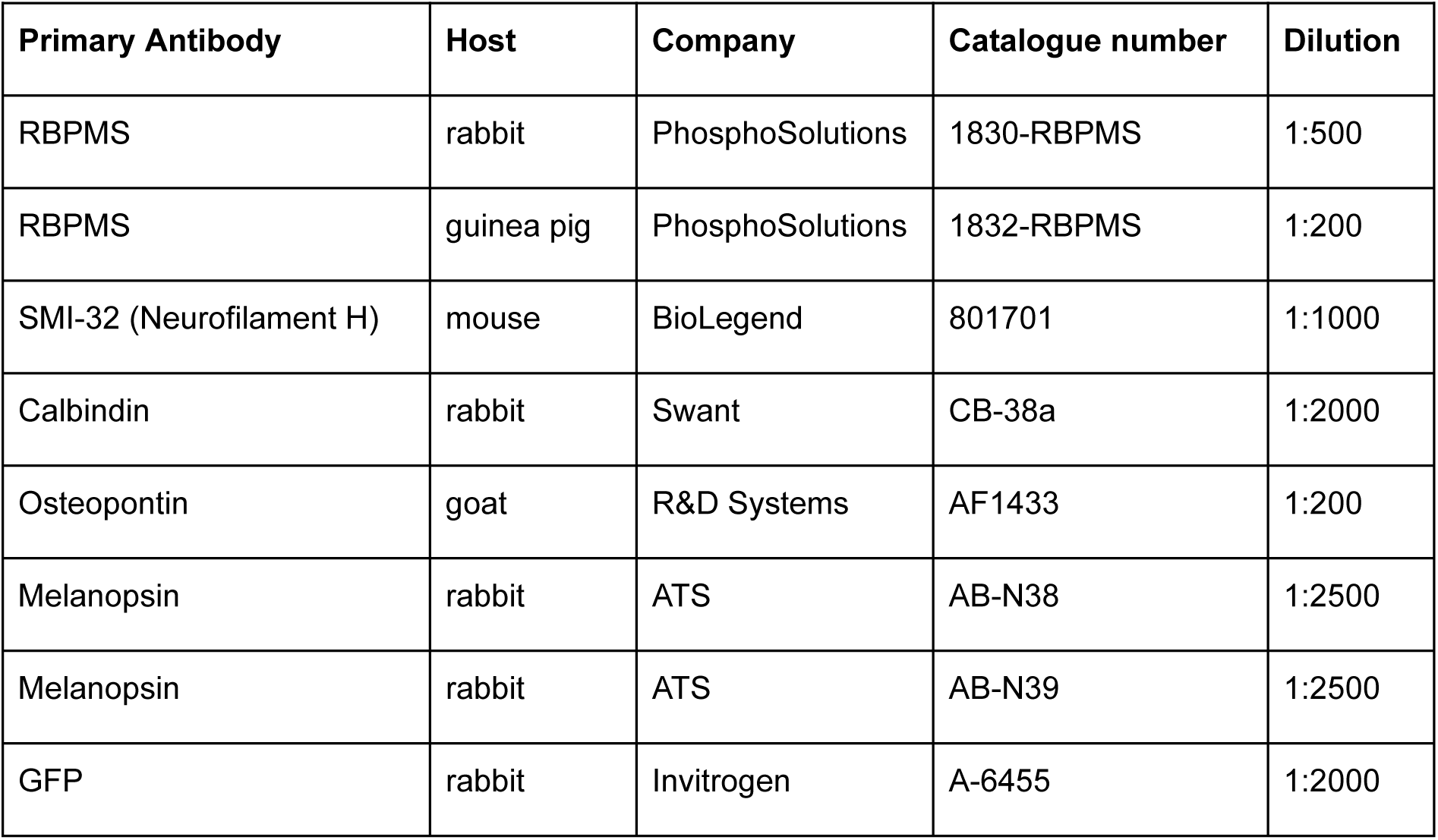
List of primary antibodies.

The tissue was mounted on glass slides and coverslipped with Vectashield (Vector Laboratories). Spacers between glass slides and coverslips were used to avoid squeezing the tissue.

### Light microscopy and image analysis

High-resolution image stacks were taken with confocal laser scanning microscopes (Leica TC SP8 and Leica TCS SL) with 40x/1.25 oil immersion objectives and z-axis increments of 0.25-0.5 µm or with a Zeiss AxioObserver equipped with an Apotome2 and a 20x/0.8 air objective and z-axis increments of 0.54 µm. Maximum intensity side view- or z-projections were done in Fiji (Schindelin et al., 2012). Brightness and contrast of the final images were adjusted in Adobe Photoshop.

Dendritic trees were traced through image stacks in Amira (Thermo Scientific) to reveal the dendritic morphology and to analyze the spatial arrangement of displaced cells in the INL relative to their counterparts in the GCL. Dendritic tracings were plotted in Matlab. DAPI labeling was aligned with the side-view projection of the dendrite tracings relative to the position of the immunolabeled dendritic trees in CorelDRAW (Corel). The inner plexiform layer (IPL) borders were determined by DAPI labeling and are represented as gray lines in figures 3, 6, and 7.

Cell bodies labeled with RBPMS, SMI-32 and calbindin or melanopsin were manually outlined and the corresponding area was measured in Fiji to calculate their diameter. For the calculation of density recovery profiles (Rodieck, 1991) and for large-scale quantification of cell densities, tile scans of image stacks from whole retinas were taken with a Leica DM6 B epifluorescence microscope equipped with a motorized stage and a 20x/0.5 air objective. Individual image stacks were automatically stitched together in the microscope software (LAS X, Leica). Next, the centers of individual cell bodies were manually marked using the *CellCounter* plugin in Fiji.

### Reconstruction of retinal wholemounts

A number of landmarks that were used in the spherical reconstruction and density calculations were marked in Fiji and Matlab. These include the outline of the flattened retina along with additional marks for cuts and tears within the said outline, the optic disc, areas where the retina ruptured internally, areas where tissue was missing, as well as prominent anatomical features such as blood vessels. The outline was estimated in rare cases where tissue was folded or missing. The R package Retistruct (Sterratt et al., 2013) was then used to reconstruct the dissected retina into the partial sphere formed by the retina in the eye. The complete retina was estimated to form a spherical cap with a rim angle of 110° (Sterratt et al., 2013). The vascularisation pattern was used to assess the quality of the reconstruction. Orientations of different retinas were aligned based on the incision along the nasal choroid fissure. The average nasal choroid fissure incision line and the average optic disc is shown in Figure 4C. The center of the optic disc was ventrally offset from the pole of the cap with an angle of 5.2±2.6°. The radius of the retinal cap, in particular of the ganglion cell layer, was set to 1.5 mm as measured in cryosections of complete mouse eyes. This is in accordance with reports from (Schmucker and Schaeffel, 2004). We discarded the radius estimated by equalizing the area of the cloverleaf and spherical cap (1.3±0.0 mm, n=10). The radius difference was likely caused by an underestimation of the area of the flattened whole-mounted retina due to potentially compressed tissue in the center of the cloverleaf around the optic nerve head while unfolding the retina during the mounting procedure.

The local cell density (i.e. density of cell body markers) was calculated in spherical coordinates. The density at a given position was the number of markers in the local neighborhood within a 10 degree radius. This radius corresponds to an arc length of 261.8 µm and a counting disc area of 0.21 mm^2^. Regions such as the optic disc or regions with insufficiently stained or damaged tissue were excluded from contributing to the area. The average density across different retinas (Figure 7) was calculated at a fixed regular grid. Reconstructed points were visualized as azimuthal equal distance projection (Figure 7).

### Matching of anatomically and electrophysiologically identified cells

Retinas were recorded as described previously (Field et al., 2007). Briefly, a small piece of isolated retina from the dorso-temporal quadrant of the eye was mounted, RGC side down, on a high-density MEA (3Brain). Recordings were analyzed offline to isolate the spikes of different cells. Candidate spike events were detected using a threshold on each electrode. The voltage waveforms on the electrode and neighboring electrodes around the time of the spike were extracted. Clusters of similar spike waveforms were identified as candidate neurons if they exhibited a refractory period. Duplicate recordings of the same cell were identified by temporal cross-correlation and removed. A monochrome binary spatial white noise was displayed on a CRT monitor at a refresh rate of 120 Hz and a stimulus pixel width of 49 µm. The grid on which stimulus pixels were presented was randomly shifted, effectively doubling the spatial resolution. Photopic light levels at a mean intensity of 2.9 mW/m^2^ were used to characterize the response properties of the recorded cells. The receptive field was approximated by the spike-triggered average. RGCs were functionally classified into cell types based on their spatiotemporal receptive field properties and spike autocorrelation function given that they formed a regular mosaic. sONɑ RGCs were identified based on their sustained responses, large receptive fields, and response pattern to full field frequency- and amplitude-modulated sweeps (Baden et al. 2016). Receptive field outlines were drawn at the 1 s.d. contour of two-dimensional Gaussian fits. The electrical image was calculated as the average voltage recorded in a 5.1 ms time window surrounding the spikes of a cell (Litke et al., 2004).

After the MEA recordings, a manual tile scan was performed while the retina was still mounted on the MEA. Images were obtained with a Leica DMLFS epifluorescence microscope equipped with a 10x air objective and stitched together in Fiji using the pairwise stitching plugin. Subsequently, the tissue was carefully mounted on a black nitrocellulose filter membrane (Millipore) with the RGC layer up and processed for immunostainings as described above. Mosaics of sONɑ RGCs were identified independently by immunolabeling and functional analyses as described above. GFP-labeled sONɑ RGCs were traced with Amira (Thermo Fisher Scientific) in the epifluorescence image from the retina when it was still mounted on the MEA to match GFP-labeled cells and the readily visible electrodes together with the corresponding electrical image. A unique mapping of anatomical and electrophysiological data was achieved by the comparison of the cell body and axon positions with the corresponding voltage deflections in the electrical images among all recorded cells.

Unless stated otherwise, all data are reported as mean ± s.d. (standard deviation).

## Results

Mouse retinas were labeled against melanopsin in combination with further immunomarkers to investigate the distribution of regularly placed ipRGCs and displaced cells. The melanopsin immunolabeling revealed a non-uniform staining pattern across the mouse retina as shown previously (Aranda and Schmidt, 2021; Hughes et al., 2013). In the dorsal retina, cell bodies and dendrites of M1 and M2 ipRGCs were clearly visible with a rather homogeneous melanopsin staining level. In contrast, the melanopsin staining pattern in the ventral retina was less homogeneous, generally weaker in most cells, but with strongly labeled (M1) cells that stood out (Fig. 1). Thus, further markers were required to reliably identify cell types and their specific distribution patterns across all retinal regions.

**Figure 1:**
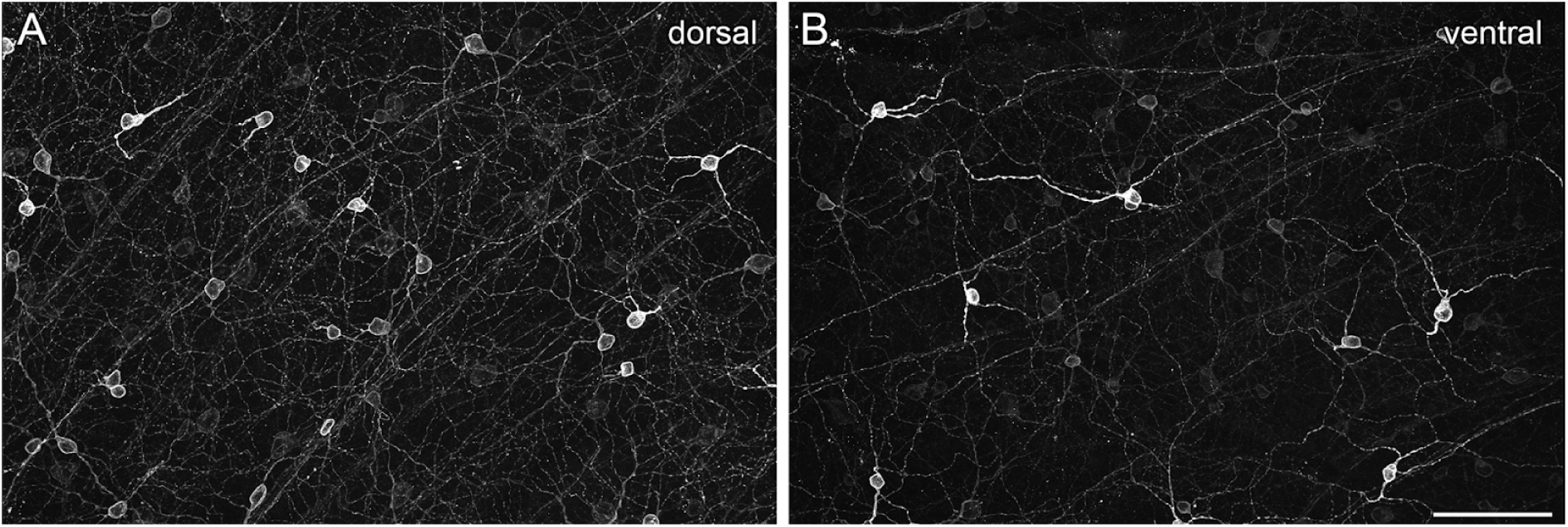
Non-uniform staining pattern of melanopsin across the mouse retina. Maximum intensity projections of structured illumination epifluorescence images labeled with melanopsin were taken from a dorsal **(A)** and a ventral **(B)** region of the same retina. Scale bar: 50 µm

### Osteopontin labels M2 ipRGCs

M1 and M2 ipRGCs are strongly immunopositive for melanopsin but a specific marker to clearly separate these two types is missing. The cytosolic protein osteopontin (Duan, 2015) is commonly used as an alpha RGC marker in mice, but it has been suggested that another type of melanopsin-containing RGCs is immunoreactive against osteopontin as well (Krieger et al., 2017; Tran et al., 2019). Thus, we analyzed the staining pattern of osteopontin in combination with melanopsin and further cell marker antibodies to determine if these combinations were suitable for a reliable separation of the RGC types of interest (Fig. 2A-D, encircled). We used SMI-32 and calbindin (Bleckert et al., 2014; Straznicky et al., 1992) to rule out a confusion with sONɑ RGCs (Figure 2 E-L, encircled), that express osteopontin and low levels of melanopsin (Estevez et al., 2012; Schmidt et al., 2014) and own observations, Fig. 2). In addition, RBPMS (RNA-binding protein with multiple splicing; (Rodriguez et al., 2014)), a general ganglion cell body immunomarker was applied to further elucidate general anatomical properties of RGC bodies. sONɑ RGCs had extraordinarily large and polygonal-shaped cell bodies (Fig. 2F, G, soma diameter 20.7±1.3 µm; n=75; see also Fig 3G).

**Figure 2:**
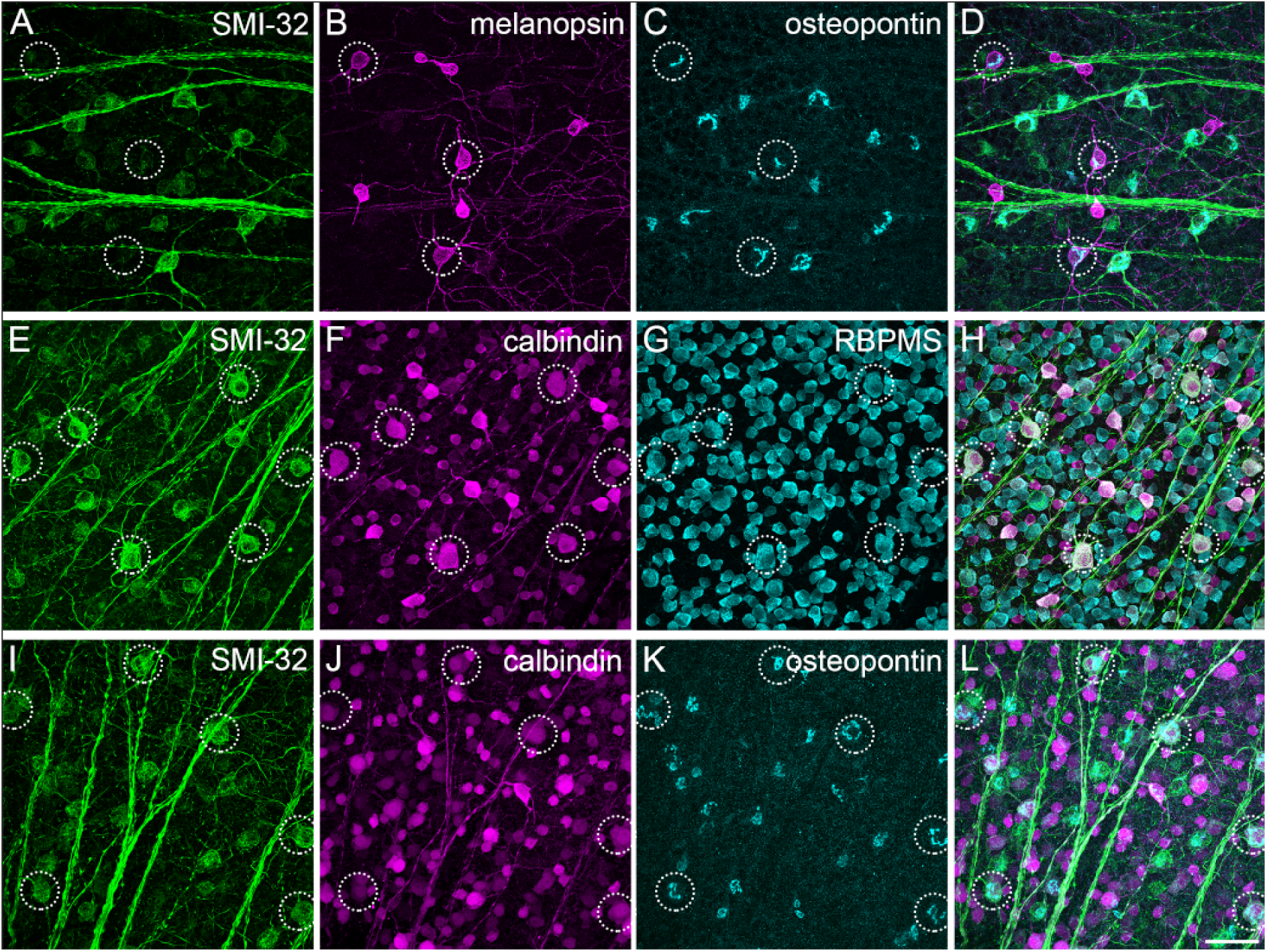
Immunomarkers for ipRGC / sONɑ cell type classification. Maximum intensity projection of confocal image stacks of the mouse GCL triple-labeled against the neurofilament marker SMI-32, melanopsin, calcium-binding protein (CaBP), RNA binding protein with multiple splicing (RBPMS), and osteopontin in different combinations. **A-D:** Cell bodies of several melanopsin-positive ipRGCs are immunonegative for SMI-32 but labeled with osteopontin (dashed circles). **E-H:** Cell bodies of sONɑ RGCs (dashed circles) are co-labeled with SMI-32, CaBP, and RBPMS. **I-L:** As E-H but with osteopontin instead of RBPMS. Scale bar: 50 µm

The marker combination presented here allowed the unequivocal classification of sONɑ RGCs as the latter were very large, strongly positive for SMI-32, and only weakly positive for melanopsin. Thus, at least one other RGC type was also immunoreactive for both osteopontin and melanopsin. To identify this type, we used dendritic stratification to classify RGCs co-labeled with both osteopontin and high levels of melanopsin. The dendrites of M1 or M2 ipRGCs stratify in sublamina S1 or S5 of the inner plexiform layer (IPL), respectively (Schmidt and Kofuji, 2009). If the immunoreactivity against osteopontin in melanopsin-labeled ipRGCs was a type-specific feature, this should be revealed by distinct stratification patterns of the corresponding cells. Thus, we analyzed the dendritic stratification of osteopontin-positive and -negative ipRGCs, which were strongly melanopsin-positive. These cells were manually traced through microscopic image stacks of immunolabeled wholemounts (Figure 3).

**Figure 3:**
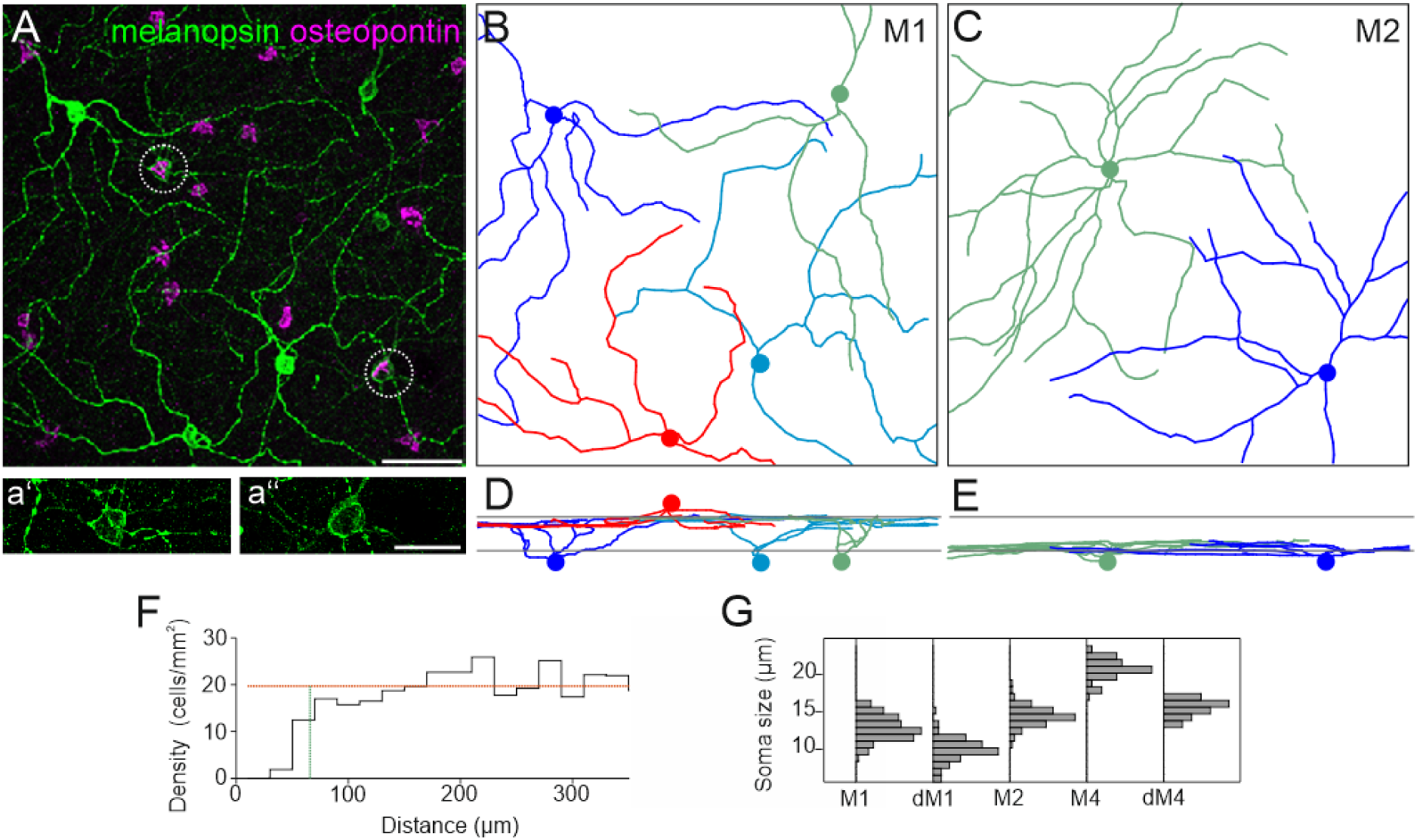
Tiling pattern and stratification levels of osteopontin-negative (M1) and -positive (M2) ipRGCs. **A:** Maximum intensity projection of a confocal image stack of wholemounted retina labeled against melanopsin and osteopontin. Dendrites of M1 and M2 ipRGCs were manually traced through the image stack in Amira. **aʻ** and **aʻʻ** show the staining pattern of melanopsin labeled cell bodies from A (dashed circles) with increased intensity. **B:** Ganglion cell skeletons of M1 ipRGCs. Colored discs indicate the positions of the cell bodies. The red skeleton at the bottom of B belongs to a displaced ganglion cell. **C:** Ganglion cell skeletons of M2 ipRGCs. **D, E:** vertical views of B, C. Gray lines indicate the inner and outer border of the IPL as determined by DAPI staining (not shown, see material and methods). **F:** Density recovery profile of M2 cells (n=281) analyzed in a wholemounted retina shows the distribution of distances between M2 ipRGCs. Effective radius of exclusion (vertical green line): 65.8 µm, cell density average over the larger half of the bins (horizontal orange line): 19.71 cells/mm^2^, bin width 20 µm. **G:** Histogram of ipRGC body diameters, M1 n=114; dM1 n=81; M2 n=81; sONɑ n=75; displaced sONɑ n=21 (all peaks normalized to equal amplitude). Scale bars: 50 µm in A, applies to A-E; 25 µm in a’’.

Cells that were negative for osteopontin but strongly positive for melanopsin exclusively stratified in the most distal part of the IPL (sublamina S1, Fig. 3D). These cells had sparsely branched dendritic arbors and relatively small cell bodies (Fig. 3G, soma diameter 12.8±1.6 µm; n=114). We identified these cells as M1 ipRGCs based on staining intensity, overall morphology, and stratification level. In contrast, osteopontin-positive, strongly melanopsin labeled ipRGCs exclusively stratified in the most proximal part of the IPL (sublamina S5, Fig. 3E). Cell bodies of these cells were slightly larger compared to M1 ipRGCs (soma diameter 14.4±1.3 µm; n=81). They had symmetric dendritic trees and more branchpoints than M1 ipRGCs, similar to the morphology that was described for M2 ipRGCs (Berson et al., 2010).

All M2 candidate cell body positions (n=281) were manually marked in an immunolabeled retina and a density recovery profile was calculated. An exclusion zone with an effective radius of 65.8 µm became apparent (Fig. 3F). This exclusion zone further supported the notion that these cells formed a single type which tiled the retina (see also Fig. 4M-O).

**Figure 4:**
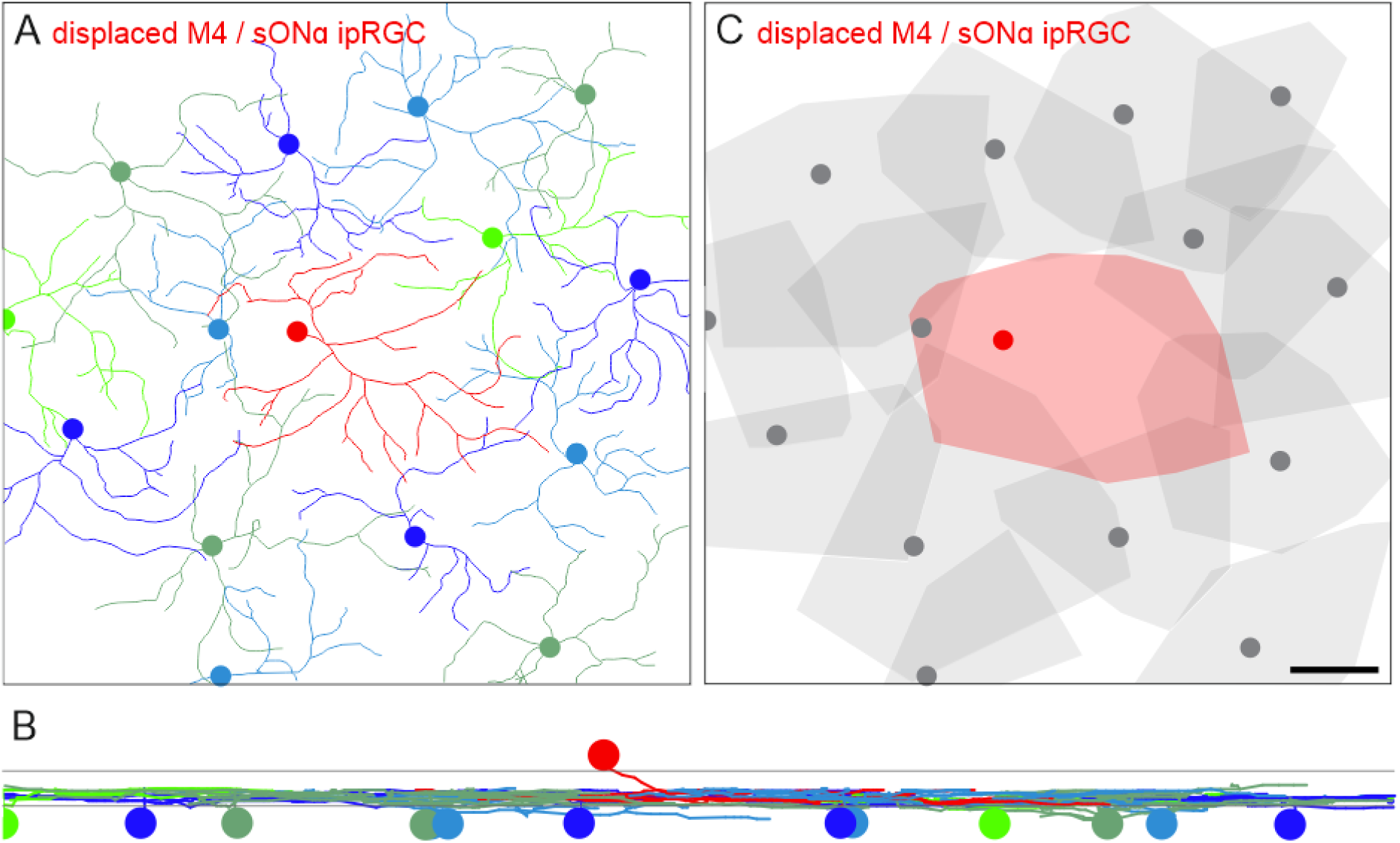
Tiling pattern and stratification levels of M4/sONɑ RGCs: Dendrites of sONɑ RGCs were manually traced through a confocal image stack of a mouse retina labeled against SMI-32 and DAPI**. A:** sONɑ RGC skeletons. Colored discs indicate the positions of the cell bodies. The red skeleton in the center belongs to a displaced sONɑ RGC. **B:** Vertical view of the RGC skeletons in A. Gray lines indicate the inner and outer border of the inner plexiform layer as determined by DAPI staining. **C:** Convex hulls encompass the skeletons as in A. The displaced cell is shown in red. Note that dendritic trees of some ganglion cells in the periphery of the image stack were not fully captured. Scale bar in C: 50 µm, also applies to A.

Together, these results showed that strongly melanopsin labeled cells, which were osteopontin-positive and SMI32-negative, represented a distinct cell type, namely M2 ipRGCs.

### Dendritic tiling pattern of regular and displaced RGC types

We analyzed the spatial arrangement of dendritic processes of displaced RGCs and their regular counterparts in the GCL to understand the dendritic organization relative to each other. For this, dendritic trees of displaced RGC types and those of their regularly placed neighbors of the same type were manually traced through confocal image stacks (Fig. 4).

The red dendritic skeleton originates from a displaced sONɑ RGC (Fig. 4A, B). The cell bodies of all remaining cells were located in the GCL. All of these cells stratified at the same level in the ON layer of the IPL. Displaced sONɑ RGCs (n=3) completed a hole in the mosaic of their regularly placed partners, and together they evenly tiled the retina (Fig. 4C).

To ensure that this dendritic tiling pattern of displaced cells is not exclusively related to sONɑ RGCs, we also analyzed the putative integration of displaced M1 ipRGC dendrites within the mosaic of regularly placed M1 ipRGCs in a ventral mouse retina (Fig. 5).

**Figure 5:**
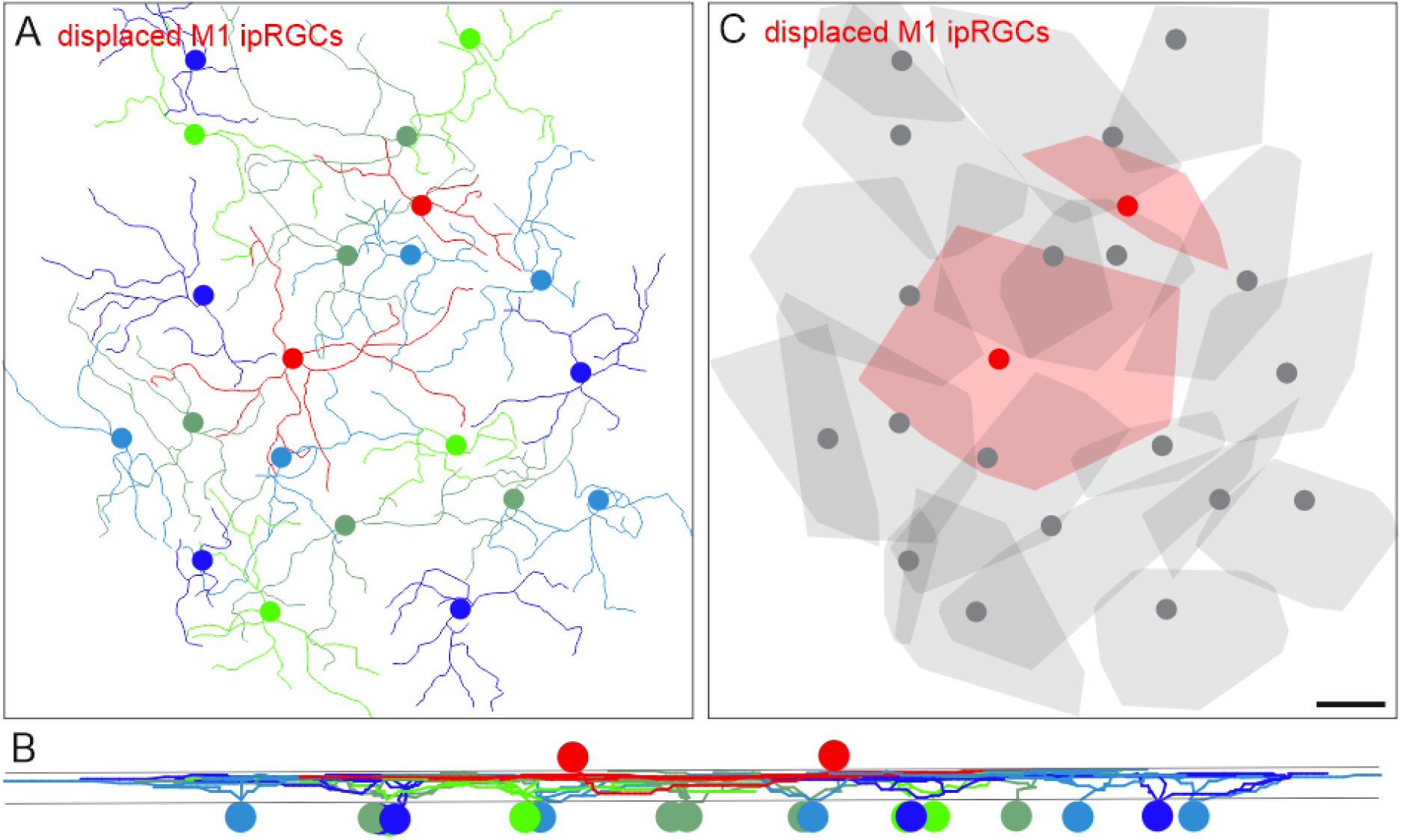
Tiling pattern and stratification levels of M1 ipRGCs. **A:** Melanopsin immunostaining was used to trace the dendrites of M1 cells through the image stack in Amira. The resulting M1 ipRGC skeletons are shown. The red skeletons represent tracings of displaced M1 cells. **B:** Vertical view of the ganglion cell skeletons in A. **C:** Convex hulls encompass the skeletons as in A. Displaced cells are shown in red. Note that dendritic trees of some ganglion cells in the periphery of the image stack were not fully captured. Scale bar in C: 100 µm, also applies to A.

The red skeletons in the center of Figure 5A indicate displaced M1 ipRGCs with cell bodies located in the INL. These cells completed the mosaic of the remaining, regular counterparts with their cell bodies located in the ganglion cell layer. All M1 ipRGCs stratified at the same level in the distal OFF layer of the IPL (Fig. 5B). As shown above for sONɑ RGCs, displaced M1 ipRGCs completed the mosaic of their regularly placed partners of the same type (Fig. 5C).

### Receptive field mosaic of regular and displaced sONɑ RGCs

We presented anatomical evidence that displaced RGCs tile the retina in concert with their regularly placed counterparts of the same type to form complete dendritic mosaics. This spatial pattern should be reflected in a homogeneous, uninterrupted mosaic of the functional receptive fields when a displaced ganglion cell is present, assuming that displaced RGCs would not differ from regular cells in terms of their light response properties.

To test this hypothesis, the receptive field organization of sONɑ RGCs was characterized by MEA recordings of spikes from a GFP-O mouse retina in response to a random noise light stimulus (Fig. 6). GFP-O retinas were used to identify GFP-positive sONɑ RGCs (Neumann et al., 2016). We recorded from the dorso-temporal retina to increase chances of recording from a displaced sONɑ RGC (see Fig. 7V-X) among regular RGCs.

**Figure 6:**
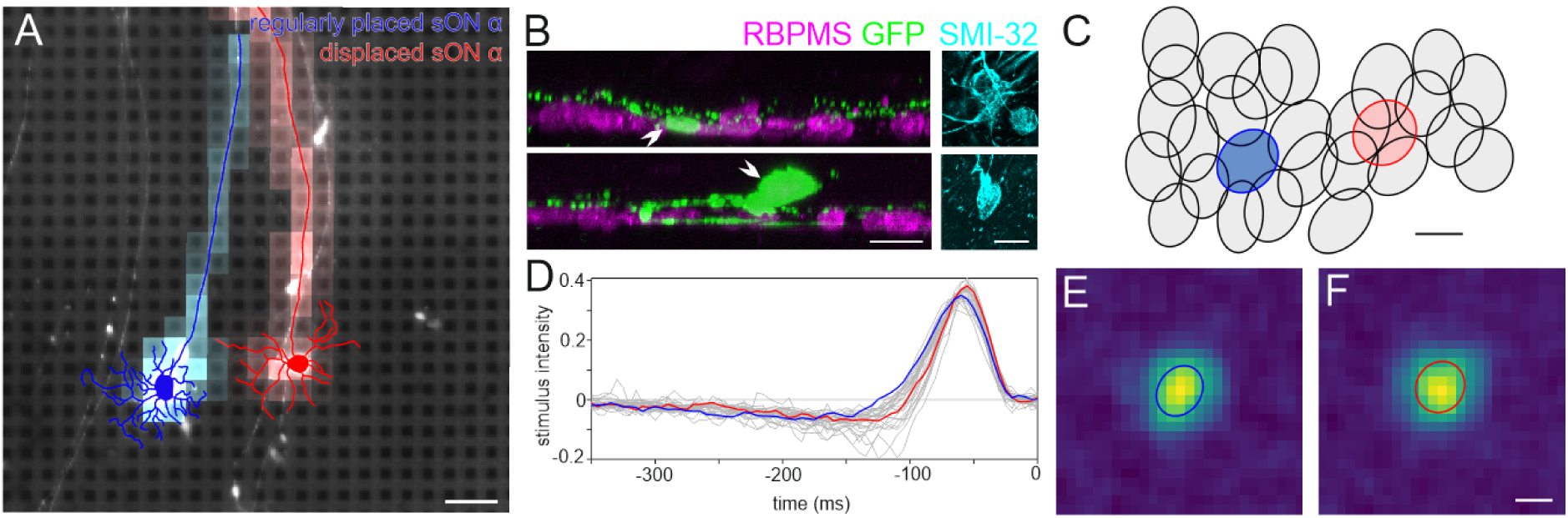
Displaced and regular sONɑ RGCs share similar light response properties. **A:** Epifluorescence image of the recorded GFP-O retina on the MEA shows native GFP fluorescence in a regularly placed sONɑ (traced in blue) and a displaced sONɑ RGC (traced in red), overlayed with the respective electrical image. Scale bar 100 µm. **B:** Projections of confocal image stacks of the same cells from A after staining with antibodies against RBPMS, GFP, and SMI-32. Left panels: Side view projections of cell bodies (arrowheads) and dendrites of regularly placed sONɑ (top) and displaced sONɑ RGC (bottom). Right panels: Top (wholemount) view projection of SMI-32 staining of cell bodies from the marked cells in the left panels (and from A). Both cells are strongly immunopositive for SMI-32. Scale bars 20 µm. **C:** Spike-triggered average stimulus time courses of all sONɑ RGCs shown in D. Blue and red lines indicate the time courses of the regularly placed and displaced RGC from A, respectively. **D**: Receptive field outlines of the sONɑ RGC mosaic. Blue and red ellipses indicate the receptive field fits of the regularly placed and displaced sONɑ RGCs in A. Scale bar 100 µm. **E:** Spatial receptive field of the regularly placed sONɑ RGC shown in A, D. **F:** Spatial receptive field of the displaced sONɑ RGC shown in A, D. Scale bar 100 µm.

**Figure 7:**
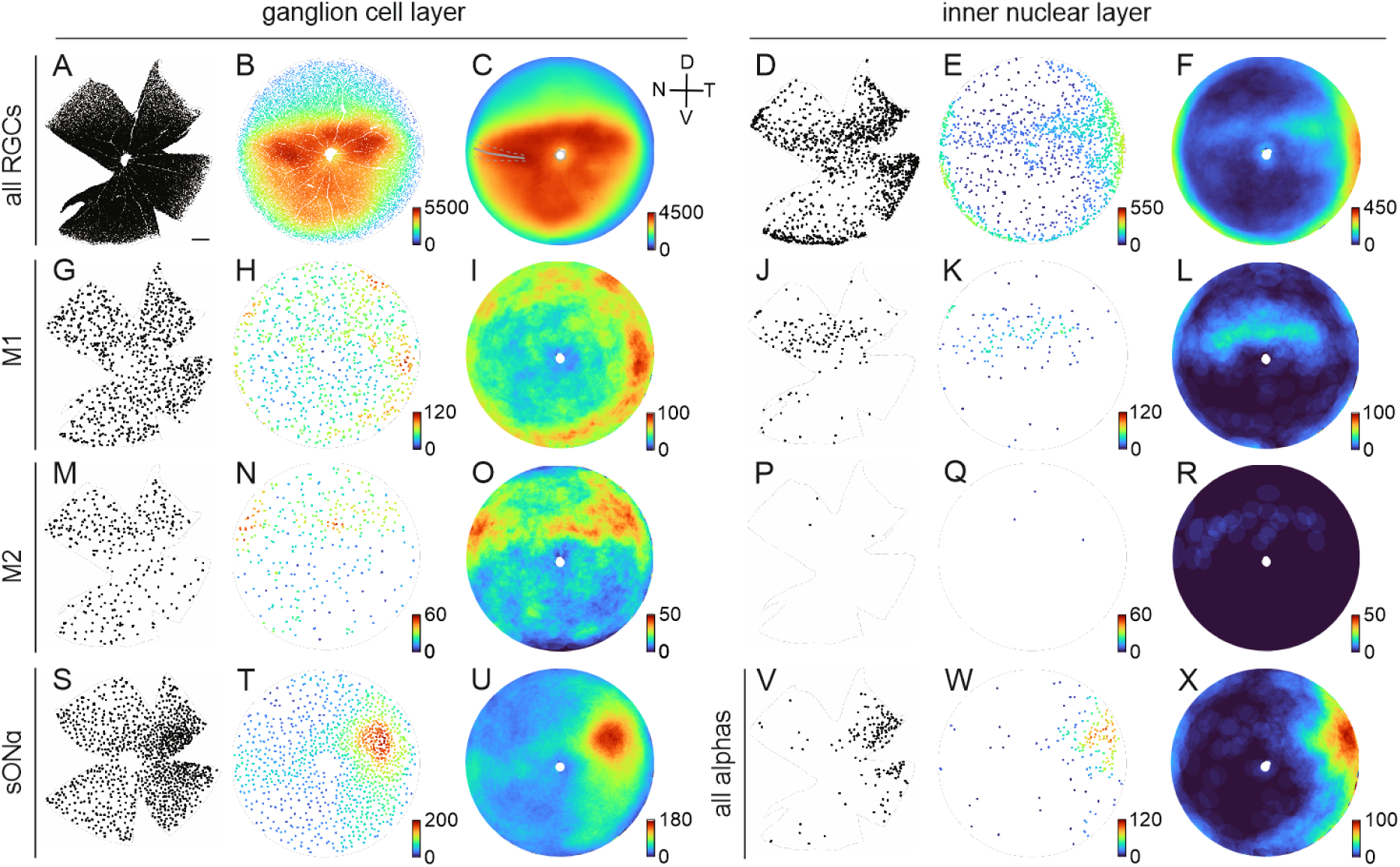
Population densities across the retina. **A:** Each dot indicates the position of an individual RGC with its cell body located in the ganglion cell layer of a retina (gray outlines) from a left mouse eye, labeled with the RGC marker RBPMS. Scale bar in A: 500 µm, applies to all cloverleaf-shaped samples. **B:** Azimuthal equal-distance projection of the reconstructed retinal sphere. The color of the dots represents the local density within a radius of 10 deg (spherical distance of ∼262 µm) of the corresponding cell. Color bar: density of cells/mm^2^. **C:** Density distributions as in B, averaged across 4 retinas. For the averages across multiple retinas, the retina orientation was adjusted based on the individual incisions along the nasal choroid fissure. Solid gray line: average fissure incision for all retinas used for analysis (n=10); dashed line: standard deviation. (D=dorsal, V=ventral, N=nasal, T=temporal) **D-F** as A-C for displaced RGCs. **G-L** as A-F for M1 ipRGCs (melanopsin positive, osteopontin negative), averaged across 5 retinas. **M-R** as A-F for M2 ipRGCs (melanopsin positive, osteopontin positive), averaged across 5 retinas. **S-U** as A-C for sONɑ RGCs in the GCL (SMI-32 positive, calbindin positive, osteopontin positive) and **V-X** as D-F for all ɑRGCs in the INL (melanopsin negative, osteopontin positive), averaged across 4 retinas. All alpha RGCs instead of sONɑ were analyzed for technical reasons, see Results for further explanations.

In one of the experiments, two nearby GFP-positive RGCs were recorded simultaneously (Fig. 6A), which were subsequently confirmed as a regularly placed sONɑ (traced in blue) and a displaced sONɑ RGC (traced in red), based on dendritic tree morphology, strong SMI-32 labeling, and the locations of cell bodies in GCL and INL, respectively (Fig. 6A,B). These two anatomically identified cells were then matched to the functional dataset from the MEA recording, by screening all candidate electrical images of all RGCs recorded in the corresponding area. We found matching pairs for both cells (Fig. 6A) based on soma positions and axon trajectories. The cells were part of the sONɑ functional mosaic (Fig. 6C) which was identified independently (see material and methods). The regularly placed and displaced sONɑ RGCs show similar receptive field properties in terms of spike response kinetics and spatial extent (Fig. 6 D-F). In line with our anatomical results, the receptive field of the displaced sONɑ RGC (red ellipse in Fig. 6C) completed the receptive field mosaic of its presumably regular-placed counterparts (gray ellipses).

### Distribution of RGC types across the retina

Here, dRGCs were observed more frequently than expected in areas other than the outer marginal zone (Dräger and Olsen, 1981). Therefore, we investigated the general distribution patterns of dRGCs relative to regular RGCs across complete retinas and in a type-specific manner by a large-scale analysis of the abovementioned immunomarkers.

We applied these marker combinations for a localization analysis of RGC bodies in immunolabeled wholemounts (Fig. 7). Stress-relief cuts to flatten the tissue yielded the typical cloverleaf-like shape of the samples. One of these cuts was placed exactly along the nasal choroid fissure during dissection for orientation of the retina. Each dot in Figure 7 indicates the position of a cell body (Fig. 7, black dots in left columns of GCL and INL panels). We reconstructed the cloverleaf shape of the tissue back into the original, almost hemispherical structure of the eyecup with Retistruct to calculate the densities of cells across the complete retinas (Sterratt et al., 2013). The local density in 3D was projected into an azimuthal equal distance plot and is indicated by the color of the dots (Fig. 7, colored dots in middle columns of GCL and INL panels). Density calculation in the reconstructed retinas allowed us to observe the averaged data of 4 or 5 retinas per analysis type (Fig. 7, colored discs in the right columns of GCL and INL panels).

### Distribution of RGC types across the retina - GCL

We identified 51946±3165 RGC bodies throughout the GCL of complete retinas (n=4), which was largely consistent with previous reports (Dräger and Olsen, 1981; Salinas-Navarro et al., 2009a). RGCs were not uniformly distributed throughout the mouse retina (Fig. 7A-C). Beyond a general central-to-peripheral gradient, the density distribution showed a steep increase in the dorsal half toward an area of highest RGC density above the optic disc and in the ventral retina.

The distribution of M1, M2, and sONɑ RGCs in the ganglion cell layer did not follow the overall density gradient of all RGCs and showed type-specific distribution patterns. 805±57 M1 ipRGCs per retina (n=5) were counted (Fig. 7 G-I). This represented a fraction of 1.6±0.2 % of all RGCs in the GCL. They had a higher density in the dorsal and temporal periphery. M2 ipRGCs showed areas of higher density in the dorsal retina (n=5, Fig. 7 M-O) (Hughes et al., 2013). We counted 336±43 M2 ipRGCs per retina. These were 0.7±0.1 % of all RGCs in the GCL. sONɑ RGCs had a density peak in the dorso-temporal retina (Bleckert et al., 2014) (Fig. 7S-U). We counted 1043±49 sONɑ RGCs per retina (n=4). These were 2.0±0.2 % of all RGCs in the GCL.

### Distribution of ipRGC types across the retina - INL

The cell bodies of dRGCs were located in the most proximal part of the INL. They were distinguished from amacrine cells by their larger cell bodies and RBPMS immunoreactivity. However, cell body size of dRGCs was relatively small compared to cell bodies of their regularly placed counterparts (Fig. 3G; soma diameters: M1 12.8±1.6 µm, n=114; displaced M1 10.0±1.6 µm, n=81; sONɑ 20.7±1.3 µm, n=75; displaced sONɑ 15.5±1.0 µm, n=21).

As shown above, different types of regularly placed RGC types exhibited distinct density patterns across the retina, consistent with previous work on the distribution of RGCs in the mouse retina (reviewed in Heukamp et al., 2020). dRGCs were expected to mostly occur in the far periphery of the retina (outer marginal zone, (Dräger and Olsen, 1981) in a potentially random fashion. However, the analysis of displaced RGCs revealed cell-type specific patterns of cell bodies in the INL, which followed neither the general density distribution of all RGCs, nor those of the corresponding cell type (Fig. 7, right INL panel).

On average, 1264±49 dRGCs were quantified per retina (n=4). This represents an amount of 2.4±0.2 % of all RGCs. Cell bodies of dRGCs were accumulated in the retinal periphery, but a large fraction was also observed in more central parts of the retina, specifically in a region above the optic disc and in the dorso-temporal area (Fig. 7D-F). 154±15 displaced M1 ipRGCs were counted per retina (n=5). This represents 16.1±1.9 % of all M1 cells. Displaced M1 cell bodies were primarily clustered in a streak-like region above the optic disc (Fig. 7J-L). In sharp contrast, M2 cell bodies were rarely found to be displaced. On average, only 5±4 displaced M2 ipRGCs were counted per retina (n=5; Fig. 7P-R). It was not possible to reliably quantify the number and distribution of displaced sONɑ RGCs. They were positive for SMI-32 and calbindin, like their regularly placed counterparts, but the cell bodies were smaller (Fig. 3G), the staining intensities were relatively low, and often superimposed by intense fluorescence of horizontal cells. Thus, we relied on the combination of RBPMS, melanopsin, and osteopontin to analyze the population of displaced ɑRGCs without any differentiation of the four types (206±35 displaced ɑRGCs per retina, n=4). The distribution of displaced ɑRGCs resembled the distribution of sONɑ RGCs in the GCL only to a certain extent (Fig. 7V-X). Similar to regularly placed sONɑ RGCs (Fig. 7S-U), the analysis of displaced ɑRGCs revealed a density peak region located in the dorso-temporal retina. However, the density peak of displaced ɑRGCs was located closer to the peripheral edge of the retina than the peak region of sONɑ RGCs in the GCL.

## Discussion

In this study, we introduced osteopontin as a marker for alpha RGCs and M2 ipRGCs which allowed us to analyze the type-specific distribution and spatial arrangement of dRGCs across complete retinas in direct relationship with their partners in the GCL. We show that the distribution of dRGCs does not follow the overall density distribution of cells in the GCL, in line with results from rat retinas (Nadal-Nicolás et al., 2014). Furthermore, we found that a displacement of cell bodies is type dependent, and we provide anatomical as well as electrophysiological evidence that dRGCs complete the tiling pattern of regular RGCs to form mosaics.

### Ganglion cell density gradients across the mouse retina

Originally, the general density distribution of mouse RGCs was described to be rather uniform, with a slight central-to-peripheral gradient (Dräger and Olsen, 1981; Jeon et al., 1998). With ongoing methodological progress regarding the use of molecular cell markers and functional analyses for cell type classification, it was shown that complex patterns of density gradients exist (Heukamp et al., 2020). Here, we provide further evidence for distinct density distributions of all RGCs and those of individual types. In line with previous results from mouse and rat retinas (Salinas-Navarro et al., 2009b, 2009a), the overall RGC density showed a steep increase from low density in the far dorsal retina to highest densities in a region above the optic disc and generally high densities in ventral retina. The density increase is roughly aligned with the transition zone of photoreceptor opsin expression (data not shown) and should therefore correspond to the horizon in the visual environment of the animal (Baden et al., 2013; Nadal-Nicolás et al., 2020).

The distribution patterns we observed for regular sONɑ and M1 ipRGCs also resembled previously reported results (Bleckert et al., 2014; Hughes et al., 2013; Johnson et al., 2021), where it was shown that the dorso-temporal density peak of sONɑ RGCs aligns with the binocular region of the visual field of the mouse. However, the density of M2 ipRGCs appeared different from previous reports (Berson et al., 2010; Hughes et al., 2013), i.e. about two times lower in our study. This discrepancy is likely caused by the analysis of whole retinas here versus quantification of much smaller tissue regions in other studies. Extrapolation of values from the M2 ipRGC density from the dorsal peak regions (Fig. 4O) for a whole retina, for instance, would yield numbers that matched previously published results. We can rule out that we missed cells in our quantification due to potentially weak melanopsin immunostaining, because weakly melanopsin-positive sONɑ RGCs were frequently observed in our samples without using further enhancement techniques (Estevez et al., 2012). Therefore, the observation of melanopsin-positive sONɑ RGCs served as an intrinsic control for optimal staining intensities.

### Density gradients and spatial arrangement of displaced ganglion cells

Displaced ganglion cells have been shown to exist in various vertebrate species, such as fish, birds, and mammals including human retina (e.g. (Bunt and Minckler, 1977; Chandra et al., 2019; Dacey et al., 2005; Dräger and Olsen, 1981; Hoshi and Sato, 2018; Mey and Johann, 2001; Nadal-Nicolás et al., 2014; Robson and Holländer, 1984; Salinas-Navarro et al., 2009b, 2009a; Stell and Witkovsky, 1973; Tachibana, 1978). However, it remained unclear whether displaced RGCs represent independent cell types, an odd part of the population of regular types, or just futile remnants of disordered neuronal development. Hallmark features of regular RGC types within local retinal regions are their uniform dendritic morphologies and the corresponding formation of independent mosaics, as well as uniform light response properties (Vlasits et al., 2019). While the existence of dRGCs was known, neither their spatial arrangement relative to regularly placed RGCs nor their light response properties have been explored.

Our data now shows that, at least for two identified RGC types of the mouse retina, dRGCs are part of the population of their regularly placed counterparts and that they are arranged as proper cellular components of the corresponding mosaic. The dendritic structure of dRGCs and the dendritic overlap with their neighbors was not obviously different from regularly placed RGCs. Interestingly, the frequency of their occurrence was not dictated by the densities of their regular counterparts, assuming that higher numbers of regular cells would increase the chance of a displacement to happen. Previous studies suggested that dRGCs in the mammalian retina are misplaced due to ontogenetic aberrations (Buhl and Dann, 1988). While the cause of displacement remains elusive, it is unlikely to happen randomly, as the patterns of displacement shown here occurred in a type-specific manner, consistently across multiple samples. Nevertheless, cell-type-specific gene expression levels during development may play a role in the formation of dRGC patterns. It was shown that the extracellular matrix protein glycogen synthase kinase 3 (GSK3) contributes to the spatial organization of RGCs (Kisseleff et al., 2021). The lack of one GSK3 allele led to increased numbers of dRGCs, but not by overproduction of cells as total RGC numbers were stable. It remains a matter of speculation that sONɑ and M1 RGCs may express lower levels of GSK3 during development, and that, therefore, their migration to the GCL is more frequently disturbed than in M2 RGCs, for instance. M2 cells were rarely found to be displaced, further supporting the notion of a cell-type dependent mechanism.

It has been shown in murine retinas that dRGCs are a highly diverse group that comprises cells with a large morphological variety (Buhl and Dann, 1988; Pang and Wu, 2011). The displaced alpha RGCs together with the displaced M1 and M2 ipRGCs added up to only ∼30% of all displaced RGCs which we observed (with M2 ipRGCs accounting for <1%). Morphological changes across cell types according to regional variations and density gradients (reviewed in (Heukamp et al., 2020) could have led to a slight overestimation of dRGC variety in previous studies. Nevertheless, one would still expect to find displaced cell bodies from many other RGC types beyond the three types investigated here. Whether all mouse dRGCs follow the rules which we found to apply to this group of cells remains to be elucidated.

### Do displaced ganglion cells serve special functional roles?

Our data shows that dRGCs complete the dendritic and corresponding functional receptive field mosaics of their regularly placed partners and that the basic light response properties do not differ between dRGCs and other cells of that mosaic. Therefore, we conclude that they belong to the same type and serve the same functional roles. However, the type-specific patterns of displacement, and the formation of a streak-like area by displaced M1 ipRGCs that is not apparent in their regularly placed partners, for instance, fuel the idea that a special role of mouse dRGCs remains to be discovered.

In the avian retina, a subpopulation of dRGCs is indeed thought to be an independent cell type (Haverkamp et al., 2021). dRGCs project exclusively to the nucleus of the basal optic root (nBOR) and are therefore thought to be responsible for the optokinetic nystagmus and involved in retinal image stabilization (Cook and Podugolnikova, 2001; Dann and Buhl, 1987; Fite et al., 1981; Simpson, 1984). The mammalian homolog of the avian nBOR is the medial terminal nucleus (MTN). It was shown that this region receives little or no input from dRGCs in rabbit (Oyster et al., 1980). However, an increased number of dRGCs in genetically modified mice was accompanied by higher numbers of RGC axon terminals in MTN (Kisseleff et al., 2021). Therefore, results regarding the innervation of the mammalian MTN from dRGCs remain inconclusive and require further studies to allow insights into potential roles of displaced RGC types in eye movements.

It has been shown that displaced ipRGCs are accumulated in an area above the optic disc in mouse retinas (Valiente-Soriano et al., 2014). Here, we provide evidence that this phenomenon occurs due to a streak-like density peak of displaced M1 ipRGCs. A conclusive classification of M1 ipRGCs remains a challenge, as they may comprise subtypes, which exhibit functional differences, distinct gene expression patterns and project to different target areas (Hattar et al., 2006; Jain et al., 2012; Li and Schmidt, 2018; Schmidt and Kofuji, 2009). It is tempting to speculate that a specific M1 subtype is responsible for the enrichment of dRGCs in this streak-like area, but this remains a subject of future studies once reliable tools are available to specifically distinguish putative M1 subtypes (Laboissonniere et al., 2019).

How could dRGCs potentially benefit from a targeted, cell-type specific displacement of their cell bodies? Changes in the common synaptic circuitry can now be ruled out, as all dendritic trees stratify at the same level and would presumably interact with the same synaptic partners. However, direct synaptic inputs to the cell bodies of dRGCs may play a key role, formed by presynaptic cells stratifying in the most distal IPL layer. An example of such a circuitry at the IPL-to-INL border is the perinuclear nests which are formed by dopaminergic amacrine cells around the cell bodies of postsynaptic AII amacrine cells (Voigt and Wassle, 1987; Casini et al., 1995; Contini and Raviola, 2003). Dendro-somatic synaptic interactions were also shown to occur at regular ganglion cell bodies with great functional impact (Grimes et al., 2022) and could represent a circuit motif for amacrine-dRGCs interactions in the INL as well. Furthermore, it is known from a subset of dRGCs to express distinct voltage-gated calcium channel subunits in murine retinas, which were not found in regular RGCs, suggesting that those dRGCs may have distinct functional properties (De Sevilla Müller et al., 2013).

Thus, our data suggests that mouse dRGCs primarily fulfill the same functional role like their regular counterparts. However, additional functional features may be discovered by a detailed study of physiological and genetic properties of dRGCs in direct association with their regularly placed partners.

## Author contributions

The study was designed by CP and MG. SD carried out immunohistochemical experiments, large-scale identification of cell types, and dendritic tracings. DRP performed multi-electrode array recordings. SD, CTB, DRP, and YA analyzed data. SD wrote the first draft of the manuscript. SD, CTB, and DRP prepared the figures. All authors edited and commented on the manuscript.

## Acknowledgements

We would like to thank Bettina Kewitz for excellent technical assistance and Alina Klaiber for assistance with the quantification of cell body positions. We also thank Karin Dedek for valuable comments on an earlier version of the manuscript. This work was supported by the VolkswagenStiftung (MG).

## References

Aranda, M.L., Schmidt, T.M. (2021). Diversity of intrinsically photosensitive retinal ganglion cells: circuits and functions. Cell Mol Life Sci CMLS 78, 889–907.

Baden, T., Berens, P., Franke, K., Román Rosón, M., Bethge, M., Euler, T. (2016). The functional diversity of retinal ganglion cells in the mouse. Nature 529, 345–350.

Baden, T., Schubert, T., Chang, L., Wei, T., Zaichuk, M., Wissinger, B., Euler, T. (2013). A Tale of Two Retinal Domains: Near-Optimal Sampling of Achromatic Contrasts in Natural Scenes through Asymmetric Photoreceptor Distribution. Neuron 80, 1206–1217.

Berry, M.H., Moldavan, M., Garrett, T., Meadows, M., Cravetchi, O., White, E., Leffler, J., von Gersdorff, H., Wright, K.M., Allen, C.N., Sivyer, B. (2023). A melanopsin ganglion cell subtype forms a dorsal retinal mosaic projecting to the supraoptic nucleus. Nat Commun 14, 1492.

Berson, D.M., Castrucci, A.M., Provencio, I. (2010). Morphology and mosaics of melanopsin?expressing retinal ganglion cell types in mice. J Comp Neurol 18.

Berson, D.M., Dunn, F.A., Takao, M. (2002). Phototransduction by Retinal Ganglion Cells That Set the Circadian Clock. 295, 5.

Bleckert, A., Schwartz, G.W., Turner, M.H., Rieke, F., Wong, R.O.L. (2014). Visual Space Is Represented by Nonmatching Topographies of Distinct Mouse Retinal Ganglion Cell Types. Curr Biol 24, 310–315.

Buhl, E.H., Dann, J.F. (1988). Morphological diversity of displaced retinal ganglion cells in the rat: A lucifer yellow study. J Comp Neurol 269, 210–218.

Bunt, A.H., Minckler, D.S. (1977). Displaced ganglion cells in the retina of the monkey. Invest Ophthalmol Vis Sci 16, 95–98.

Casini, G., Rickman, D.W., Brecha, N.C. (1995). AII amacrine cell population in the rabbit retina: identification by parvalbumin immunoreactivity. J Comp Neurol 356, 132–142.

Chandra, A.J., Lee, S.C.S., Grünert, U. (2019). Melanopsin and calbindin immunoreactivity in the inner retina of humans and marmosets. Vis Neurosci 36, E009.

Collin, S.P. (2008). A web-based archive for topographic maps of retinal cell distribution in vertebrates. Clin Exp Optom 91, 85–95.

Contini, M., Raviola, E. (2003). GABAergic synapses made by a retinal dopaminergic neuron. Proc Natl Acad Sci 100, 1358–1363.

Cook, J.E., Podugolnikova, T.A. (2001). Evidence for spatial regularity among retinal ganglion cells that project to the accessory optic system in a frog, a reptile, a bird, and a mammal. Vis Neurosci 18, 289–297.

Curcio, C.A., Allen, K.A. (1990). Topography of ganglion cells in human retina. J Comp Neurol 300, 5–25.

Dacey, D.M., Liao, H.-W., Peterson, B.B., Robinson, F.R., Smith, V.C., Pokorny, J., Yau, K.-W., Gamlin, P.D. (2005). Melanopsin-expressing ganglion cells in primate retina signal colour and irradiance and project to the LGN. Nature 433, 749–754.

Dann, J.F., Buhl, E.H. (1987). Retinal ganglion cells projecting to the accessory optic system in the rat. J Comp Neurol 262, 141–158.

De Sevilla Müller, L.P., Liu, J., Solomon, A., Rodriguez, A., Brecha, N.C. (2013). Expression of Voltage-Gated Calcium Channel α2δ4 Subunits in the Mouse and Rat Retina. J Comp Neurol 521, 2486–2501.

Doi, M., Uji, Y., Yamamura, H. (1995). Morphological classification of retinal ganglion cells in mice. J Comp Neurol 356, 368–386.

Dräger, U.C., Olsen, J.F. (1981). Ganglion cell distribution in the retina of the mouse. 20, 9.

Duan, X. (2015). Subtype-Specific Regeneration of Retinal Ganglion Cells following Axotomy: Effects of Osteopontin and mTOR Signaling. 22.

Ecker, J.L. (2010). Melanopsin-Expressing Retinal Ganglion-Cell Photoreceptors: Cellular Diversity and Role in Pattern Vision. 12.

Estevez, M.E., Fogerson, P.M., Ilardi, M.C., Borghuis, B.G., Chan, E., Weng, S., Auferkorte, O.N., Demb, J.B., Berson, D.M. (2012). Form and Function of the M4 Cell, an Intrinsically Photosensitive Retinal Ganglion Cell Type Contributing to Geniculocortical Vision. J Neurosci 32, 13608–13620.

Feng, G., Mellor, R.H., Bernstein, M., Keller-Peck, C., Nguyen, Q.T., Wallace, M., Nerbonne, J.M., Lichtman, J.W., Sanes, J.R. (2000). Imaging Neuronal Subsets in Transgenic Mice Expressing Multiple Spectral Variants of GFP. Neuron 28, 41–51.

Field, G.D., Sher, A., Gauthier, J.L., Greschner, M., Shlens, J., Litke, A.M., Chichilnisky, E.J. (2007). Spatial Properties and Functional Organization of Small Bistratified Ganglion Cells in Primate Retina. J Neurosci 27, 13261–13272.

Fite, K.V., Brecha, N., Karten, H.J., Hunt, S.P. (1981). Displaced ganglion cells and the accessory optic system of pigeon. J Comp Neurol 195, 279–288.

Grimes, W.N., Sedlacek, M., Musgrove, M., Nath, A., Tian, H., Hoon, M., Rieke, F., Singer, J.H., Diamond, J.S. (2022). Dendro-somatic synaptic inputs to ganglion cells contradict receptive field and connectivity conventions in the mammalian retina. Curr Biol 32, 315–328.e4.

Hattar, S., Kumar, M., Park, A., Tong, P., Tung, J., Yau, K.-W., Berson, D.M. (2006). Central projections of melanopsin-expressing retinal ganglion cells in the mouse. J Comp Neurol 497, 326–349.

Hattar, S., Liao, H.-W., Takao, M., Berson, D.M., Yau, K.-W. (2002). Melanopsin-Containing Retinal Ganglion Cells: Architecture, Projections, and Intrinsic Photosensitivity. Science 295, 1065–1070.

Haverkamp, S., Albert, L., Balaji, V., Němec, P., Dedek, K. (2021). Expression of cell markers and transcription factors in the avian retina compared with that in the marmoset retina. J Comp Neurol 529, 3171–3193.

Haverkamp, S., Mietsch, M., Briggman, K.L. (2022). Developmental errors in the common marmoset retina. Front Neuroanat 16.

Heukamp, A.S., Warwick, R.A., Rivlin-Etzion, M. (2020). Topographic Variations in Retinal Encoding of Visual Space. Annu Rev Vis Sci 6, 237–259.

Hoshi, H., Sato, F. (2018). The morphological characterization of orientation-biased displaced large-field ganglion cells in the central part of goldfish retina. J Comp Neurol 526, 243–261.

Hughes, S., Watson, T.S., Foster, R.G., Peirson, S.N., Hankins, M.W. (2013). Nonuniform Distribution and Spectral Tuning of Photosensitive Retinal Ganglion Cells of the Mouse Retina. Curr Biol 23, 1696–1701.

Jain, V., Ravindran, E., Dhingra, N.K. (2012). Differential expression of Brn3 transcription factors in intrinsically photosensitive retinal ganglion cells in mouse. J Comp Neurol 520, 742–755.

Jeon, C.-J., Strettoi, E., Masland, R.H. (1998). The Major Cell Populations of the Mouse Retina. J Neurosci 18, 8936–8946.

Johnson, K.P., Fitzpatrick, M.J., Zhao, L., Wang, B., McCracken, S., Williams, P.R., Kerschensteiner, D. (2021). Cell-type-specific binocular vision guides predation in mice. Neuron 109, 1527–1539.e4.

Kisseleff, E., Vigouroux, R.J., Hottin, C., Lourdel, S., Thomas, L., Shah, P., Chédotal, A., Perron, M., Swaroop, A., Roger, J.E. (2021). Glycogen Synthase Kinase 3 Regulates the Genesis of Displaced Retinal Ganglion Cells3. ENeuro 8, ENEURO.0171-21.2021.

Krieger, B., Qiao, M., Rousso, D.L., Sanes, J.R., Meister, M. (2017). Four alpha ganglion cell types in mouse retina: Function, structure, and molecular signatures. PLOS ONE 12, e0180091.

Laboissonniere, L.A., Goetz, J.J., Martin, G.M., Bi, R., Lund, T.J.S., Ellson, L., Lynch, M.R., Mooney, B., Wickham, H., Liu, P., Schwartz, G.W., Trimarchi, J.M. (2019). Molecular signatures of retinal ganglion cells revealed through single cell profiling. Sci Rep 9, 15778.

Li, J.Y., Schmidt, T.M. (2018). Divergent projection patterns of M1 ipRGC subtypes. J Comp Neurol 526, 2010–2018.

Litke, A.M., Bezayiff, N., Chichilnisky, E.J., Cunningham, W., Dabrowski, W., Grillo, A.A., Grivich, M., Grybos, P., Hottowy, P., Kachiguine, S., Kalmar, R.S., Mathieson, K., Petrusca, D., Rahman, M., Sher, A. (2004). What does the eye tell the brain?: Development of a system for the large-scale recording of retinal output activity. IEEE Trans Nucl Sci 51, 1434–1440.

Mey, J., Johann, V. (2001). Dendrite development and target innervation of displaced retinal ganglion cells of the chick (Gallus gallus). Int J Dev Neurosci Off J Int Soc Dev Neurosci 19, 517–531.

Nadal-Nicolás, F.M., Kunze, V.P., Ball, J.M., Peng, B.T., Krishnan, A., Zhou, G., Dong, L., Li, W. (2020). True S-cones are concentrated in the ventral mouse retina and wired for color detection in the upper visual field. ELife 9, e56840.

Nadal-Nicolás, F.M., Salinas-Navarro, M., Jiménez-López, M., Sobrado-Calvo, P., Villegas-Pérez, M.P., Vidal-Sanz, M., Agudo-Barriuso, M. (2014). Displaced retinal ganglion cells in albino and pigmented rats. Front Neuroanat 8, 99.

Nasir-Ahmad, S., Lee, S.C.S., Martin, P.R., Grünert, U. (2019). Melanopsin-expressing ganglion cells in human retina: Morphology, distribution, and synaptic connections. J Comp Neurol 527, 312–327.

Neumann, S., Hüser, L., Ondreka, K., Auler, N., Haverkamp, S. (2016). Cell type-specific bipolar cell input to ganglion cells in the mouse retina. Neuroscience 316, 420–432.

Oyster, C.W., Simpson, J.I., Takahashi, E.S., Soodak, R.E. (1980). Retinal ganglion cells projecting to the rabbit accessory optic system. J Comp Neurol 190, 49–61.

Oyster, C.W., Takahashi, E.S., Hurst, D.C. (1981). Density, soma size, and regional distribution of rabbit retinal ganglion cells. J Neurosci Off J Soc Neurosci 1, 1331–1346.

Pang, J.-J., Wu, S.M. (2011). Morphology and Immunoreactivity of Retrogradely Double-Labeled Ganglion Cells in the Mouse Retina. Investig Opthalmology Vis Sci 52, 4886.

Provencio, I., Rodriguez, I.R., Jiang, G., Hayes, W.P., Moreira, E.F., Rollag, M.D. (2000). A Novel Human Opsin in the Inner Retina. J Neurosci 20, 600–605.

Quattrochi, L.E., Stabio, M.E., Kim, I., Ilardi, M.C., Michelle Fogerson, P., Leyrer, M.L., Berson, D.M. (2019). The M6 cell: A small-field bistratified photosensitive retinal ganglion cell. J Comp Neurol 527, 297–311.

Robson, J.A., Holländer, H. (1984). Displaced ganglion cells in the rabbit retina. Invest Ophthalmol Vis Sci 25, 1376–1381.

Rodieck, R.W. (1991). The density recovery profile: A method for the analysis of points in the plane applicable to retinal studies. Vis Neurosci 6, 95–111.

Rodriguez, A.R., de Sevilla Müller, L.P., Brecha, N.C. (2014). The RNA binding protein RBPMS is a selective marker of ganglion cells in the mammalian retina. J Comp Neurol 522, 1411–1443.

Rousso, D.L., Qiao, M., Kagan, R.D., Yamagata, M., Palmiter, R.D., Sanes, J.R. (2016). Two Pairs of ON and OFF Retinal Ganglion Cells Are Defined by Intersectional Patterns of Transcription Factor Expression. Cell Rep 15, 1930–1944.

Salinas-Navarro, M., Jiménez-López, M., Valiente-Soriano, F.J., Alarcón-Martínez, L., Avilés-Trigueros, M., Mayor, S., Holmes, T., Lund, R.D., Villegas-Pérez, M.P., Vidal-Sanz, M. (2009a). Retinal ganglion cell population in adult albino and pigmented mice: A computerized analysis of the entire population and its spatial distribution. Vision Res 49, 637–647.

Salinas-Navarro, M., Mayor-Torroglosa, S., Jiménez-López, M., Avilés-Trigueros, M., Holmes, T.M., Lund, R.D., Villegas-Pérez, M.P., Vidal-Sanz, M. (2009b). A computerized analysis of the entire retinal ganglion cell population and its spatial distribution in adult rats. Vision Res 49, 115–126.

Schindelin, J., Arganda-Carreras, I., Frise, E., Kaynig, V., Longair, M., Pietzsch, T., Preibisch, S., Rueden, C., Saalfeld, S., Schmid, B., Tinevez, J.-Y., White, D.J., Hartenstein, V., Eliceiri, K., Tomancak, P., Cardona, A. (2012). Fiji: an open-source platform for biological-image analysis. Nat Methods 9, 676–682.

Schmidt, T.M., Alam, N.M., Chen, S., Kofuji, P., Li, W., Prusky, G.T., Hattar, S. (2014). A Role for Melanopsin in Alpha Retinal Ganglion Cells and Contrast Detection. Neuron 82, 781–788.

Schmidt, T.M., Chen, S.-K., Hattar, S. (2011). Intrinsically photosensitive retinal ganglion cells: many subtypes, diverse functions. Trends Neurosci 34, 572–580.

Schmidt, T.M., Kofuji, P. (2009). Functional and Morphological Differences among Intrinsically Photosensitive Retinal Ganglion Cells. 7.

Schmucker, C., Schaeffel, F. (2004). A paraxial schematic eye model for the growing C57BL/6 mouse. Vision Res 44, 1857–1867.

Simpson, J.I. (1984). The accessory optic system. Annu Rev Neurosci 7, 13–41.

Stabio, M.E., Sabbah, S., Quattrochi, L.E., Ilardi, M.C., Fogerson, P.M., Leyrer, M.L., Kim, M.T., Kim, I., Schiel, M., Renna, J.M., Briggman, K.L., Berson, D.M. (2018). The M5 Cell: A Color-Opponent Intrinsically Photosensitive Retinal Ganglion Cell. Neuron 97, 150–163.e4.

Stell, W.K., Witkovsky, P. (1973). Retinal structure in the smooth dogfish, Mustelus canis: General description and light microscopy of giant ganglion cells. J Comp Neurol 148, 1–31.

Sterratt, D.C., Lyngholm, D., Willshaw, D.J., Thompson, I.D. (2013). Standard Anatomical and Visual Space for the Mouse Retina: Computational Reconstruction and Transformation of Flattened Retinae with the Retistruct Package. PLoS Comput Biol 9, e1002921.

Stone, J. (1965). A quantitative analysis of the distribution of ganglion cells in the cat’s retina. J Comp Neurol 124, 337–352.

Straznicky, C., Vickers, J.C., Gábriel, R., Costa, M. (1992). A neurofilament protein antibody selectively labels a large ganglion cell type in the human retina. Brain Res 582, 123–128.

Sun, W., Li, N., He, S. (2002). Large-scale morphological survey of mouse retinal ganglion cells. J Comp Neurol 451, 115–126.

Tachibana, M. (1978). Displaced ganglion cells in carp retina revealed by the horseradish peroxidase technique. Neurosci Lett 9, 153–157.

Tran, N.M., Shekhar, K., Whitney, I.E., Jacobi, A., Benhar, I., Hong, G., Yan, W., Adiconis, X., Arnold, M.E., Lee, J.M., Levin, J.Z., Lin, D., Wang, C., Lieber, C.M., Regev, A., He, Z., Sanes, J.R. (2019). Single-Cell Profiles of Retinal Ganglion Cells Differing in Resilience to Injury Reveal Neuroprotective Genes. Neuron 104, 1039–1055.e12.

Valiente-Soriano, F.J., García-Ayuso, D., Ortín-Martínez, A., Jiménez-López, M., Galindo-Romero, C., Villegas-Pérez, M.P., Agudo-Barriuso, M., Vugler, A.A., Vidal-Sanz, M. (2014). Distribution of melanopsin positive neurons in pigmented and albino mice: evidence for melanopsin interneurons in the mouse retina. Front Neuroanat 8, 131.

Vlasits, A.L., Euler, T., Franke, K. (2019). Function first: classifying cell types and circuits of the retina. Curr Opin Neurobiol 56, 8–15.

Voigt, T., Wassle, H. (1987). Dopaminergic innervation of A II amacrine cells in mammalian retina. J Neurosci 7, 4115–4128.

Wässle, H., Peichl, L., Boycott, B.B. (1981). Dendritic territories of cat retinal ganglion cells. Nature 292, 344–345.

Wässle, H., Riemann, H.J. (1978). The Mosaic of Nerve Cells in the Mammalian Retina. 24.

Zhang, Y., Kim, I.-J., Sanes, J.R., Meister, M. (2012). The most numerous ganglion cell type of the mouse retina is a selective feature detector. Proc Natl Acad Sci U S A 109, E2391–E2398.

